# Perception of Speech Turn Dynamics is preserved in Congenitally Deaf children with Cochlear Implants

**DOI:** 10.1101/2023.05.22.538554

**Authors:** Céline Hidalgo, Christelle Zielinski, Sophie Chen, Stéphane Roman, Eric Truy, Daniele Schön

## Abstract

Perceptual and speech production abilities of children with cochlear implants (CI) are usually tested by word and sentence repetition or naming tests. However, in their daily life they show very heterogeneous language skills. Here, we describe a way of increasing the ecological validity of language assessment, promoting the use of close to real-life listening situations. The setup consists in watching the audio-visual conversation of two individuals. Children’s gaze-switches from one speaker to the other serve as a proxy of their prediction abilities. Moreover, to better understand the basis and the impact of anticipatory behaviour, we also measured children’s ability to understand the dialogue content, their speech perception and memory skills as well as their rhythmic skills. Importantly, we compared children with CI performances with those of an age-matched group of children with NH. While children with CI revealed poorer speech perception and verbal working memory abilities than NH children, there was no difference in gaze anticipatory behaviour. Interestingly, in children with CI only, we found a significant correlation between dialogue comprehension, perceptive skills and gaze anticipatory behaviour. Our results confirm and extend to a conversational context previous findings showing an absence of predictive deficits in children with CI. The current design seems an interesting avenue to provide an accurate and objective estimate of anticipatory language behaviour in a rather ecological conversational context also with young children.

## Introduction

Even when early implanted, children with cochlear implants (CI) show heterogeneous language skills (van Wieringen & Wouters, 2015). Indeed, although they are mostly included in mainstream schools and can, sometimes, achieve language scores close to those of children with normal-hearing (NH) (Hayes et al., 2009), several children with CI still show deficits in vocabulary knowledge (Lund, 2016) and morphological, syntactic or narrative skills (Boons et al., 2013). The best predictors of such variability in speech and language levels of children with CI seem to be the age of implantation, the child’s residual hearing and the parents socio-economic status (Niparko et al., 2010; Nicholas & Geers, 2018; see Ruben, 2018 for a review). Nonetheless, these factors do not explain the entire heterogeneity found in their language outcomes (Houston, 2022).

This may be partly linked to the lack of ecological validity of the tests. Indeed, it has been shown that adding ecologically valid variables can increase the variance accounted for (Chaytor et al., 2006) and it is surely one of the great challenges of hearing research in the years to come (Keidser et al., 2020). Indeed, perceptual and speech production abilities of children with CI are usually tested by word and sentence repetition or naming tests (see e.g. Yoshinaga-Itano et al., 2010 for language assessment in a longitudinal study or Dettman et al., 2016 for a multi-centre study) but are not integrated into the ecological contexts they encounter in their daily lives. Thus, it is important to increase the ecological validity of language assessment and promote the use of perceptual and linguistic assessments close to real-life listening situations (Lunner et al., 2020).

Adopting more ecologically valid tests may also improve our understanding of language deficits insofar as different processes may be at work in speech perception tests compared to a real situation such as listening to continuous speech. These multiple processes, that go well beyond auditory processing, may be even more complex in an interactive communication context (Carlile & Keidser, 2020). Indeed, if understanding a narrative requires more complex processing than understanding words or sentences out of context (Chambers & Juan, 2008, Xu et al., 2005), in conversational contexts, comprehension is even less of a passive process but would involve, on the contrary, prediction/anticipation capacities at different levels of representations (Friston et al., 2020, Pickering & Gambi, 2018). These predictions would be particularly solicited during degraded listening conditions of speech (Sohoglu & Davis, 2020; Strauß et al., 2013) and linked to the processing of the global comprehension of a conversation between two individuals (Hidalgo et al., 2022).

One way to directly measure predictions and their impact on comprehension was proposed by Grisoni and colleagues (2021), but requires recording EEG during sentence processing as well as controlled linguistic material. Recently, we suggested a new approach inspired by the work of Casillas and Frank (2017), more ecologically amenable, consisting in the measure, while watching audio-visual conversations of two individuals, of gaze switches, a temporally-resolved behavioural proxy of internal predictions. Interestingly we found a relation between gaze switches and global understanding of the dialogue content, in particular when speech is strongly degraded (Hidalgo et al., 2022).

Furthermore, temporal predictions play a major role in other cognitive domains, as for instance music. Importantly, temporal predictions may share common processes in speech and music since both build on hierarchical temporal structures that engage similar and partially overlapping cognitive and neural mechanisms (Schön & Morillon, 2018, Patel & Morgan, 2017). In children with CI, we recently showed a link between rhythmic and language perceptive abilities. Not only children with CI suffer from rhythmic deficits compared to their NH peers in simple sensori-motor tasks (like following the beat of music), but the better they can reproduce rhythms with a complex hierarchical structure, the better they perceive language (Hidalgo et al., 2021).

The aim of this study was twofold. First, to assess the temporal predictive abilities of children with CI in a near-ecological speech perception context: listening to a conversation and anticipating turns. Second, to analyse the link between temporal predictions, comprehension and rhythmic abilities. To compare children with CI and NH predictive abilities and their links with language processing during ecological conditions of speech perception, we have used a recently developed paradigm (Hidalgo et al., 2022) wherein children fitted with an eye-tracker, watch videos of two individuals talking to each other. Gaze-switches from one speaker to the other serve as a proxy of children’s prediction abilities. At the end of each dialogue, children have to answer questions about the content of the dialogues. Finally, we collected data of sensori-motor skills using musical rhythmic tasks with different levels of complexity as well as speech perception and working memory tests.

We make the hypothesis that 1) children with CI will have poorer temporal predictive skills compared to children with NH during conversation processing, 2) their predictive abilities will be related to their comprehension and their complex sensori-motor skills.

## Materials and Methods

### Participants

Two groups of children, aged between 8 and 12 years (mean = 10.5 years; SD = 1.3 months) were selected. In the experimental group, 19 children with cochlear implants were recruited from two ENT paediatric French centers: one in Lyon, (Edouard Herriot Hospital) and one in Marseille, (La Timone Hospital). All children (nine girls) fitted with cochlear implants were French native speakers and suffered from severe to profound hearing loss without additional disorders. These children were mostly fitted with bilateral cochlear implants (seventeen). Two of them wore only one CI without a contralateral hearing aid. All children were included in mainstream schools (see demographics for more details in Supplementary Table 1). We screened all children for their ability to understand the content of videos showing two individuals having a conversation (similar to the experimental material). In the control group, fifteen children with normal hearing were recruited in Lyon and Marseille. These children (eleven girls) were all French native speakers without any known visual, speech or cognitive disorders. This group of children passed an hearing screening at 20dB HL at 250 Hz, 500Hz, 1000 Hz, 2000 Hz, 4000 Hz and 8000 Hz using a custom screening hearing test made in Python 3.7.9 (Expyriment version 0.10.0). They were age-matched with children from the experimental group and a Wilcoxon-test between groups revealed no age difference (W = 72, *p* = 1).

Two children with normal hearing were excluded from the analysis for loss of eye-tracking data (> 50%). Three children with cochlear implants were excluded due to a difficulty in understanding the screening stimuli. Three others were excluded for loss of eye-tracking data (> 50%) and one for too many blinks in the eye-tracking data, leaving thirteen and twelve children in each group.

All children and their parents were informed of the procedure of the study, which was approved by the Sud Méditerranée V Ethics Committee (ID RCB : 2020-A02556-33).

### Procedure

#### Perceptual, cognitive and rhythmic assessments

For all children, the tests took place in a quiet room in Marseille or in Lyon hospitals in the ENT Paediatric Departments.

#### Speech perception test

Children listened to and repeated 21 non-words composed of 3 to 6 syllables (mean = 4.14 syllables, SD = 1.28) extracted from the TERMO French test (Busquet & Descourtieux, 2003). Repetitions of the stimuli were allowed when necessary. Scoring comprised an estimate of the accuracy for the global item (0 or 1, max score = 21) and the number of correct syllables reproduced, max score = 87).

#### Span tests

In order to assess the verbal working memory we used stimuli from the Wechsler Intelligence Scale for Children IV test (ECPA, 2005). Children listened to and repeated in the same order (direct span) or in the reverse order (indirect span) series of digits of increasing length. The test ended when the child made two consecutive errors at a given length. Scoring takes place at the item level (correct or incorrect). These digits were scored by the experimenter as 0 (failure) or 1 (success). The maximum score children could get for direct and indirect span was 14 and 16. Actual scores were transformed in % of correct responses with respect to the total score. The stimuli of the speech perception and memory span tests were delivered orally by the experimenter with lip-reading available.

#### Rhythmic tests

In order to assess the rhythmic abilities, children tapped as synchronously as possible along a metronome, for one minute (beat = 666ms). Then, they repeated the same task with a musical excerpt (“Chocolate” by the band The 1975 ; beat = 603ms). Finally we also assessed the ability to reproduce complex rhythms. In this task, children listened to a complex rhythm that is repeated eight times in a row. Children have to imitate the whole rhythmic pattern (and not the beat) by tapping along the rhythm. We used eight different patterns. Importantly, the use of a timbre with a short attack time and a short duration ensures a low spectral complexity of these stimuli.

Using home-made Matlab scripts, sensori-motor synchronization on the metronome and the musical beat were plotted on a polar scale in which each tap is represented on a circle of 360° by and angle relative to the expected beat time (= 0° on the circle). Taps in a trial are treated as unitary vectors; the resulting vector is calculated to quantify sensori-motor synchronization to the beat (see Sowiński & Dalla Bella, 2013). The length of this vector, named synchronization consistency (from 0 to 1), is a measure of synchronization performance (0 = uniform random distribution; 1 = perfect phase synchronization).

For the sensori-motor synchronization on complex rhythmic patterns, we considered only the last three repetitions in the scoring; the first five were considered as the learning phase. Note however that analyses were run on approximately ninety taps per child. First, we extracted the temporal series of tapping from the audio recording. To this aim we used the SciPy function find_peaks with a minimal distance of 100 ms and a prominence of 2 SD. We used an objective procedure as in Hidalgo et al., 2021, to score children’s performances. This measure is based on computing a correlation between children’s tap timing and the rhythms to be reproduced. The score of each performance was defined as the maximum value of these correlations.

Children’s speech productions (non-words, digits and sentences repetition) and tapping were recorded with a digital recorder (Zoom, Handy Recorder, H4n).

The speech from sentences and video-dialogues as well as rhythmic sounds were delivered through loudspeakers (Behringer MS 16) which were located on both sides of the screen.

### Experimental task: prediction during conversation

#### Stimuli

The videos consisted of three audiovisual stimuli of ∼4.5 minutes each, showing the head of a man and a woman having a conversation. The themes of the conversations differed and were inspired from novels and pieces of theatre for children. Dialogues were as joyful as possible to maintain children’s attention during the whole duration of the videos. Dialogues contained on average 59 turns (min = 46; max = 66). Turns lasted on average 3.7 s (SD = 2.7) and global speech rate was approximately 5Hz. 60% of these turns were questions/responses adjacency pairs (min = 56 %; max = 66%) in order to elicit predictive behaviours as suggested in Casillas & Frank, 2017 (Casillas & Frank, 2017). We used Final Cut Pro X to set the gap duration at speakers’ turn to 500ms in order to have homogeneous turn conditions and allow anticipatory gaze behaviour (cf. Foulsham et al., 2010; Casillas and Frank, 2017). Importantly visual cues such as head and eye movements as well as anticipatory mouth and lips movements were absent in the present stimuli(see Hidalgo et al., 2022 for more details on how visual cues were controlled).

### Procedure

Children were equipped with the monocular head-mounted Pupil core eye-tracker (Pupil Labs GmbH, Berlin, Germany) with their chin rested on a chinrest, fixed at the table at ∼60 cm from the screen (24’’ with a resolution of 1920 × 1080px and a refresh rate of 100Hz). Pupil Capture software (v 3.0.7) launched the eye-tracker calibration and recording. Before the calibration procedure, we ensured that QRcodes stuck on the four screen’s corners (to detect the screen surface) and children’s pupils were well detected by the eye-tracker world and eye cameras respectively. Then we run a 5 target calibration followed by a 4 target test procedure. During the task, children were instructed to listen and watch each video carefully in order to be able to answer twelve questions after each dialogue. These were close questions and could be scored as failure or success (0/1), allowing a maximum score of twelve per dialogue. The video and associated questions were delivered using the OpenSesame software (Mathôt et al., 2012).

### Stimuli and eye-tracking synchronization

A custom-made sync box based on an Arduino micro-controller ensured the synchronization between the eye-tracking dat a and the auditory signal embedded in the video file. The stereo sound went from the stimulation laptop to the sync box (audio cable). There, the stereo was split and the first channel containing the speech signals of the dialogues sent directly to the loudspeakers. The second channel signal went through the micro-controller. This channel contained an audio spike placed at the onset and at the end of each dialogue. The Arduino detected the audio trigger and sent a code to the serial port that was subsequently detected and transmitted via a Python script to the Pupil Core network API, an API which supports fast and reliable communication and real-time access to data. This ensured, for each dialogue, a precise synchronization of the eye-tracking.

### Eye-tracking data processing

Gaze data relative to the screen surface were then extracted using the Pupil Player software (v 3.0.7) and gaze switches were detected using custom Matlab scripts. In this study we used the gaze switch from one speaker to the other as a proxy of anticipatory behaviour with respect to the upcoming turn onset (see Figure 1). We considered a temporal window around the turn allowing to keep both anticipatory and non-anticipatory gaze switches, resulting in a -1/+1s window around the turn onset. Note, nonetheless that, as expected, most shifts occurred in the inter-turn gap, that is before the turn onset (Foulsham et al., 2010; Keitel et al. 2013; Casillas and Frank 2017, see Figure 1A, right panel).

**Figure 1.**
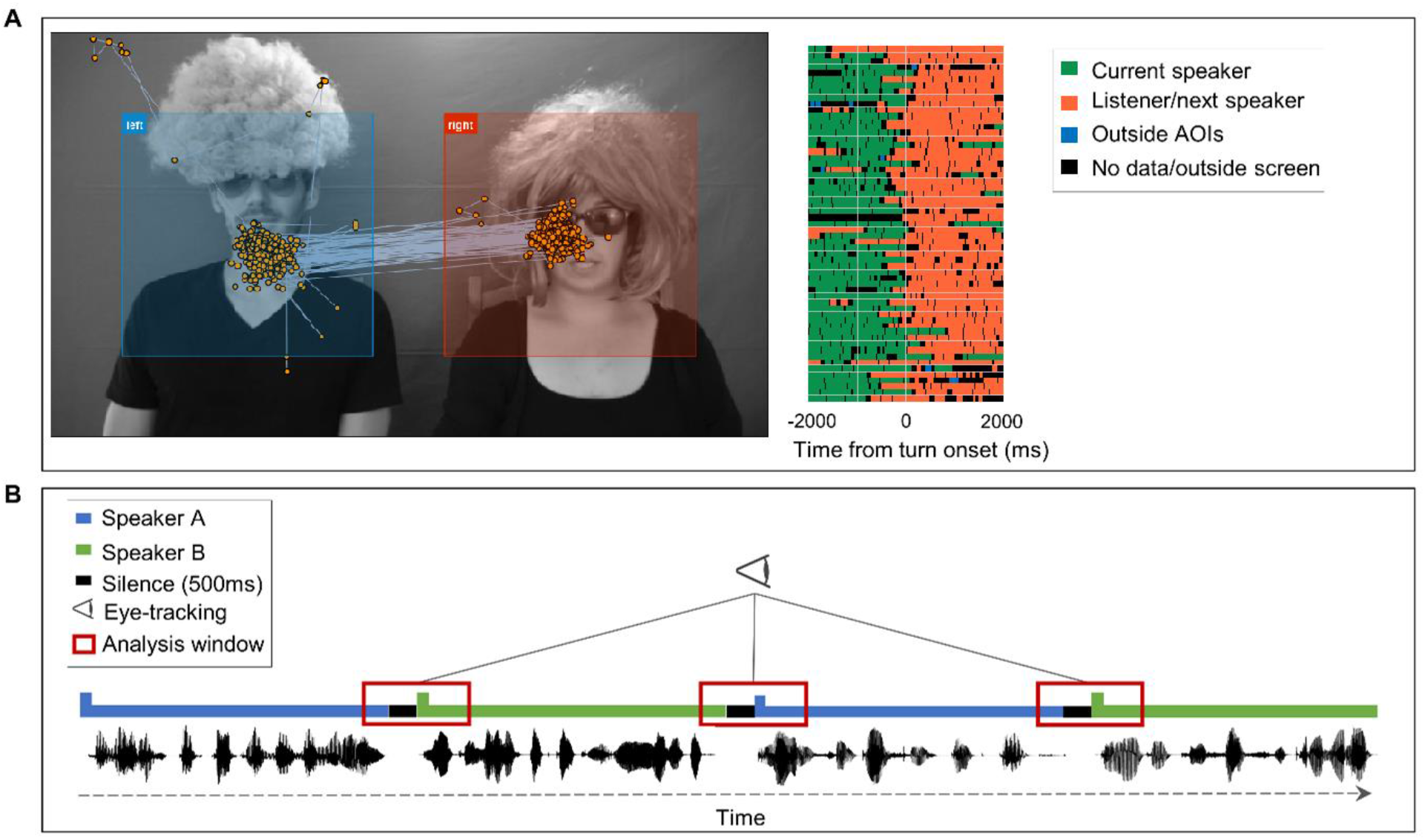
Schematic illustration of the task and analyses. A. On the left, snapshot of one video illustrating the areas of interest (AOI) and the fixation points of a single participant during the whole video duration (the photograph depicts two researchers who consented to its publication). On the right, gaze behaviour of the same participant showing the gaze switch for every turn of a single dialogue with respect to turn onset (time zero). B. Schematic illustration of the turns in the dialogue, the controlled gap between turns (500ms) and the gaze switch window of analysis.

Moreover, we also applied three supplementary criteria to filter spurious gaze switches. First, gaze before and after the switch should fall within an area of interest (AOIs), defined as a stationary rectangle surrounding each face (see Figure 1A left). Second, the gaze switch should be preceded by at least 100ms fixation on the current speaker. Third, it should be followed by at least 150ms fixation to the next speaker. Concerning the possible back and forth gaze behaviour preceding turn (∼5% in our data), we kept only the first gaze switch.

In short, we computed a gaze switch latency relative to the upcoming turn onset time. A positive value indicates thus a gaze switch after the turn onset, while a negative value indicates a gaze switch before the turn onset (Figure 1A right).

### Statistical modeling

All statistical analysis was computed using R (R Core Team, 2021) and the lme4 package (Bates et al., 2015). We computed Linear Mixed Models (LMM) with *group* as a predictor variable, to evaluate the difference of performances between children with Cochlear Implant (CI) and children with Normal-Hearing (NH) on: 1) the number of good syllables reproduced during the non-word repetition task, 2) the correlation’s coefficients obtained from the complex rhythm sequences, 3) the gaze switches during the video-dialogues. We modelled *subject* as a random effect in all analyses. We added *item* as a supplementary random effect in the model for the non-word repetition. Because the complex rhythm task may be sensitive to *musical practice* and *working memory*, we modelled these two variables as a confounding factor in the LMM.

To evaluate the differences between groups on 1) the comprehension score, 2) the direct span and 3) the indirect span measures, we computed Generalized Linear Mixed Models because of the binary nature of the variables (0 or 1), with *group* as a predictor variable and *subject* as random effect. Statistical significance of the fixed effects was assessed by model comparison using the Akaike Information Criterion, thus arbitrating between complexity and explanatory power of the models. Normality and homoscedasticity of the residuals of all the models were systematically visually inspected. Reported p values are Satterthwhaite approximations obtained with the lmerTest package (Kuznetsova et al., 2017).

For the metronome and musical beat synchronisation tests that yield one summary statistics for participant (vector length, that is the response consistency), we computed Linear Models, with *group* as a predictor variable and *musical practice* as a confounding factor.

Finally, in order to analyse the relationship between comprehension level, non-word perception skills, gaze switch latencies during videos and synchronization skills on complex rhythmic sequences, we computed Spearman’s coefficient correlations between these measures, for each group separately (12 subjects in each group). When relevant, we computed a Linear Model for the significant correlations also introducing the covariates of interest in the model. All the assumptions of the models were tested using the Global Validation of Linear Model Assumptions package and for each group comparison, we computed Cohen’d effect size.

## Results

### Perceptual, cognitive and rhythmic performances

As expected, children with CI showed lower perceptual capacities compared to children with NH. In the non-word repetition task, CI children accuracy (number of correct syllables) was lower compared to NH children (*β* = -0.476, SE = 0.0728, t = -6.549, *p* < 0.001, Cohen’s *d* = 0.436, see Fig. 2). Children with CI also showed poorer verbal memory performances than children with NH as assessed by the forward span test (*β* = -0.796, SE = 0.22, t = -3.579, *p* < 0.001, Cohen’s *d* = 0.391, see Fig. 3 panel A). Groups also differed in terms of working memory capacities, as assessed by the inverse span test, although this difference remained marginally significant (*β* = -0.384, SE = 0.207, t = -1.854 *p* = 0.063, Cohen’s *d* = 0.189, see Fig. 3 panel B). Sensori-motor synchronisation skills were also poorer in children with CI compared to those of children with NH. They tapped less consistently on the metronome beat (*β* = -0.058, SE = 0.026, t = -2.258, *p* = 0.033, Cohen’s *d* = 0.903, see Fig. 4) but this difference was no longer significant when controlling for musical practice (p = 0.227). Concerning synchronisation consistency on the musical beat, we found no difference between groups (p = 0.123) even when controlling for the musical practice (*p* = 0.45). However, children with CI’s synchronisation abilities were not as good as those of children with NH on complex rhythmic sequences (*β* = -0.079, SE = 0.025, t = -3.175, *p* = 0.004, Cohen’s *d* = 0.864, see Fig. 5) and this difference was still significant when controlling for working memory and musical practice (*β* = -0.077, SE = 0.025, t = -3.007, *p* = 0.006).

**Figure 2.**
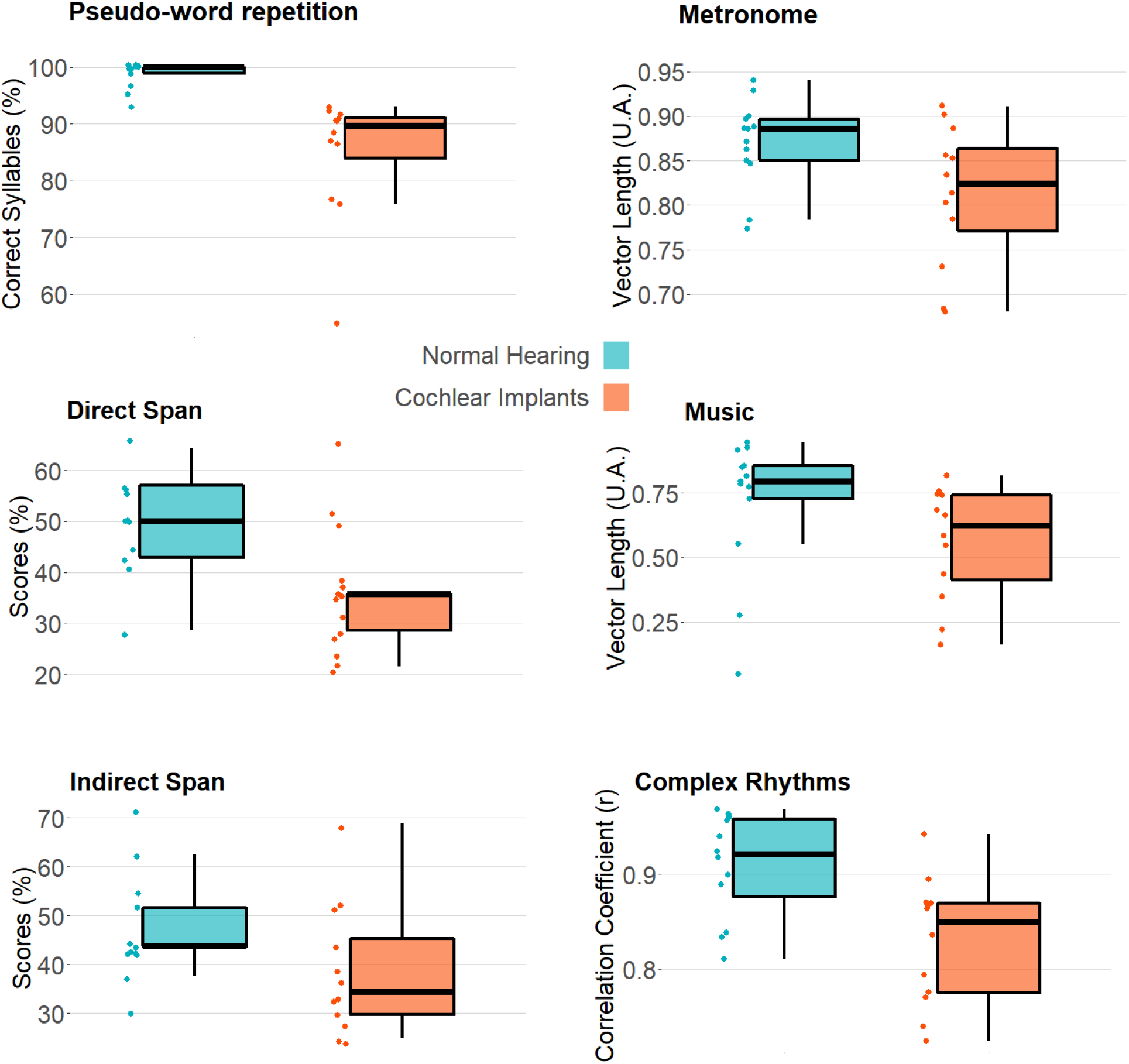
Boxplots of the results of the speech repetition, direct and indirect span, tapping to a metronome, tapping to music and complex rhythm reproduction tasks for children with normal hearing and cochlear implants.

**Figure 3.**
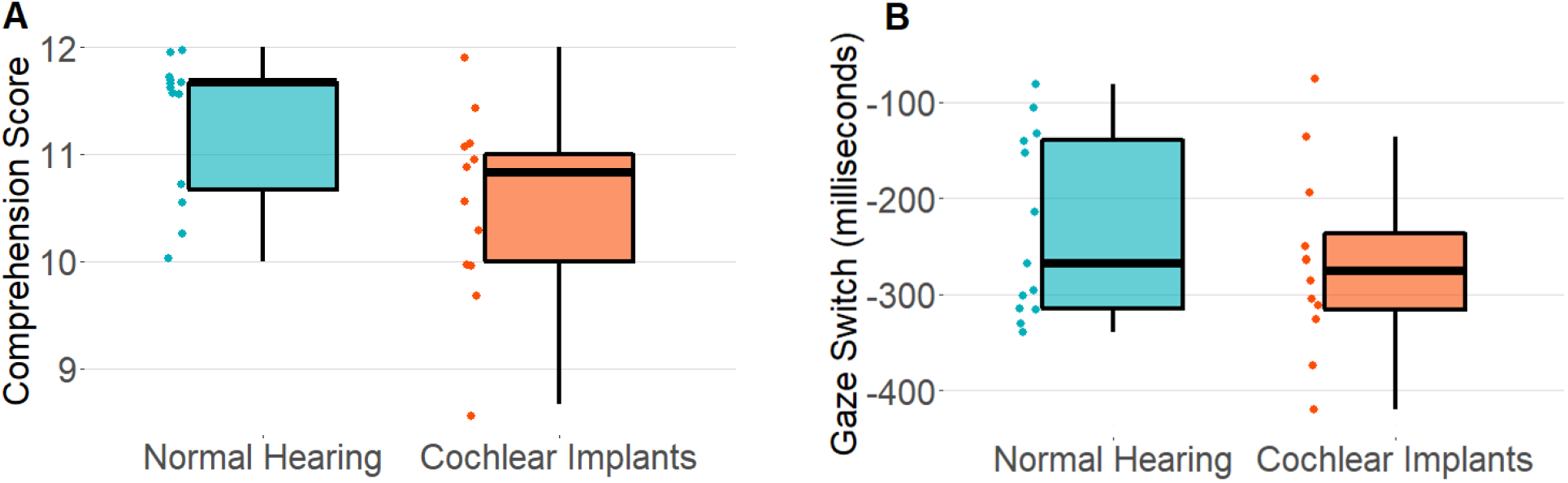
Boxplots of the results for the comprehension score and the gaze switch in the experimental task wherein children are first watching two people having a conversation. The comprehension score for each child is reported as the average of the correct answers across the three dialogues (based on twelve questions each). The gaze switch for each child is reported as the average time for gaze switch around the 177 turns (negative values indicate a gaze switch before the onset of the next speaker).

### Comprehension and temporal prediction performances

Compared to the children with NH, dialogue comprehension scores were lower for children with CI (*β* = -0.886, SE = 0.34, t = -2.606, *p* = 0.009, Cohen’s *d* = 0.231, see Fig. 6). This was despite the fact that children with CI spent more time looking at the current speakers than their NH peer (W = 274, p < 0.001). When investigating the gaze behaviour in the two groups, we found that there were no significant differences between children with CI and their NH peers, (*β* = -36.28, SE = 38.54, t = -0.941, *p* = 0.356) (see Fig.7).

#### Correlations

Correlations on separated groups revealed in children with CI a positive link between the capacities to understand dialogues and their perceptual language skills, measured by non-words repetition (*r* = 0.71, *β* = 0.197, SE = 0.061, t = 3.227, *p* = 0.009). Gaze anticipatory behaviour while watching the dialogues was also associated to good perceptual language skills (*r* = -0.65; *β* = -6.584, SE = 2.407, t = -2.736, *p* = 0.021). Gaze behaviour was also associated to comprehension scores (*r* = -0.53, *p* = 0.074), and this relation became even more apparent when taking into account memory capacities of children with CI as a co-variate (*β* = -27.63, SE =10.63, t = -2.599, *p* = 0.028). Sensorimotor synchronization abilities and in particular performance in the complex rhythm reproduction task were also associated to gaze behaviour (r=-0.39) and perceptual language skills (r=0.43) although these medium size correlations did not reach significance due to the small sample size. By contrast, we did not find any significant correlation between variables in children with NH (all *ps* > 0.05) mostly due to a ceiling effect or smaller variability of their performance in most of the skills assessed (cf Figure 2).

## Discussion

In this study, we investigated speech-turn temporal predictions abilities in congenitally deaf children with CI. To this end, we recorded eye movements and used gaze switch latencies as a proxy of anticipatory behaviour while children were watching videos of two individuals having a conversation. Moreover, in order to better understand the basis and the impact of anticipatory behaviour, we also measured children’s ability to understand the dialogue content, their speech perception and memory skills as well as their rhythmic skills. Importantly, we compared children with CI performances with those of an age-matched group of children with NH.

As expected, children with CI revealed poorer speech perception and verbal working memory abilities than NH children. They also showed poorer sensori-motor synchronisation skills in particular on complex rhythmic sequences. On the experimental task, children with CI showed poorer dialogue comprehension, spent more time looking at the current speaker but showed no differences in gaze anticipatory behaviour compared to children with NH. Interestingly, in children with CI only, we found a significant correlation between dialogue comprehension, perceptive skills and gaze anticipatory behaviour.

A first relevant element in discussing these results is that the task we used, inspired by previous work of Casillas and Frank (2017) and further refined and assessed in adults (Hidalgo et al., 2022) allows to assess both language comprehension and anticipatory verbal behaviour in a rather ecological setup. Importantly, the stimuli we used allow to interpret the findings in terms of auditory predictions. Indeed, a great deal of visual cues that typically occur in conversation, such as hand, eye, head and mouth movements, were controlled in the present study (see methods). Thus, anticipatory gaze behaviour can occur mostly based on linguistic information that here comprised both speech and lip reading inputs.

Another relevant aspect is the fact that, while a few previous studies have used indirect measures (e.g. manipulating the predictability of the context) to assess the ability of children with CI to predict words in speech (Smiljanic and Sladen, 2013; Holt et al., 2016), here we use a direct measure, eye movements, as a direct proxy of predictions in a rather ecological task. The use of gaze latencies had already been successfully employed in a visual word completion task, showing a reliable use of semantic informative cues to predict upcoming words in children with hearing loss (Holt, et al., 2021; Blomquist et al., 2021). However, the current setup, by tracking continuous gaze shift during a conversation more closely resemble to a real-life situation compared to sentence completion using a picture.

The lack of group differences concerning linguistic predictive skills goes somewhat against the idea that children with cochlear implants have difficulties extracting structure from and learning sequential input patterns, possibly due to auditory deprivation in the first years of life (Conway et al., 2009; Deocampo et al., 2018). It stands also in contrast with a reduced use of contextual information (Conway et al., 2014; Eisenberg et al., 2002; Stelmachowicz et al., 2000) and slow language processing (Burkholder & Pisoni, 2003; Pisoni et al., 2011; Holt et al., 2016) among children with hearing loss. However, recently Pesnot Lerousseau and colleagues developed a modified serial reaction time task where cochlear implanted children and normal hearing children had to react to auditory sequences that embed multiple statistical regularities. No differences between cochlear implanted children and their normal hearing peers were visible at the group level (Pesnot Lerousseau et al., 2022). These results are more in line with the present findings as they indicate that auditory statistical learning is preserved in congenitally deaf children with cochlear implants. Importantly, we extend these findings to a much more ecological context, here watching the movies of two people having a conversation wherein multiple levels of predictions are required to efficiently anticipate turns.

Our data also show that children with CI seem to rely more on the visual modality. This probably allows compensating the poorer speech integration and comprehension that may otherwise reduce their anticipatory behaviour (Bergeson and Pisoni, 2004; Peterson et al., 2010; Holt et al., 2023). Indeed, recently we showed that, using the same setup in normal hearing adults, speech degradation increases the time spent on the current speaker and reduces the anticipatory gaze behaviour (Hidalgo et al., 2022). Here, we only observed a greater time spent on the current speaker in children with CI compared to children with NH, but no differences at the level of anticipatory gaze behaviour. It is possible that children with CI extra time on the current speaker compensated for their poorer audiovisual integration and understanding of the dialogues. Indeed, improving audiovisual integration may improve syntactic and semantic processing that play an important role to anticipate turns (Lewis et al., 2017; Blomquist et al., 2021; Holt et al., 2023). Moreover, children may also rely on supra-segmental features such as coarse prosodic patterns (e.g. questions and answers) that may be partly preserved (Carter et al., 2002), although they may depend on other variables such as musical environment (Torppa et al., 2014).

While clear group differences are absent in gaze predictive behaviour, we did find that temporal predictions during conversation correlate with both speech integration abilities (non-word repetition) and dialogue comprehension. More precisely, the more the anticipatory gaze behaviour, the better speech integration and dialogue comprehension in children with CI. Interestingly, in the normal hearing adult experiment, the relation between gaze anticipatory behaviour and dialogue comprehension was maximal in the strongly speech degraded condition and almost absent in the normal speech condition (Hidalgo et al., 2022). These results under the strongly degraded and normal conditions resemble to the results we obtained with children with CI and with NH, respectively. Indeed, previous work showed that adverse listening conditions increase the uncertainty of the speech signal, thus urging listeners to exploit prior knowledge or expectations to constrain perception (Smiljanic & Sladen, 2013; Sohoglu & Davis, 2020). This possibly explains the lack of correlation between gaze anticipatory behaviour and dialogue comprehension in children with normal hearing who did not experience degraded speech in the present experiment.

Our study has a certain number of limitations. First, while our design goes indeed in the direction of ecological studies, it cannot yet be considered as a real-life listening situation and further efforts will be required in order to be able to accurately track speech perception and anticipatory behaviour in natural conditions (Lunner et al., 2020). In particular, testing in a quiet booth is not representative of real-life conditions in terms of background noise, which may explain the rather good performance in terms of comprehension and in particular in anticipatory gaze behaviour. Second, while we do reproduce previous findings showing impaired speech perception (Caldwell & Nittrouer, 2013), comprehension (Niparko et al., 2010; Caselli et al., 2012), working memory (Nittrouer et al., 2013; AuBuchon et al., 2015)) and temporal processing (Good et al., 2017; Polonenko et al., 2017; Hidalgo et al., 2021), our paradigm does not allow to understand how children with CI manage to have a normal anticipatory gaze behaviour for turn taking. As stated above they may more heavily rely upon audiovisual integration, coarse prosodic patterns or lexico-semantic or syntactic structures. Further studies may manipulate for instance the prosodic content of conversational speech, such as intonative turn continuation contour which are finer prosodic cues (Portes and Bertrand, 2005), to see to what extent it is important for cochlear implant wearer during turn processing. Nonetheless, considering the absence of visual cues that typically play a role in turn taking (Auer, 2022), the fact that children with CI have a similar anticipatory gaze behaviour than children with NH is a very good news and an interesting starting point to further understand the difficulties they encounter in conversational context (Crowe & Dammeyer, 2021).

To conclude, our results confirm and extend to a conversational context previous findings showing an absence of predictive deficits in children with CI (Hall et al., 2018; von Koss Torkildsen et al., 2018; Holt et al., 2021, Blomquist et al., 2021, Pesnot Lerousseau et al., 2022). Moreover, the present design seems an interesting avenue to provide an accurate and objective estimates of anticipatory behaviour in a rather ecological conversational context (Keidser et al., 2020). Also note that this type of design has been previously used with very young children, showing that they spontaneously made anticipatory gaze switches by age two and continue improving through age six (Casillas & Frank, 2017). Thus, it may be possible to use it to refine diagnosis at a rather early age and soon after cochlear implantation.

Finally the link between anticipatory gaze behaviour and perceptual language skills and speech comprehension seem to indicate that improving anticipatory behaviour may be beneficial for children with CI. Moreover, simpler anticipatory behaviour such as the one required for rhythm perception and production seem to be linked to perceptual language skills and speech comprehension (see also Hidalgo et al., 2021). Establishing a link between temporal prediction abilities and level of comprehension in an ecological conversational setup may allow to refine the speech therapy care techniques, as for instance by favouring rhythmic music stimulation (Hidalgo et al., 2017, 2019).

## Author contributions

Conceptualization C.H. and D.S.; Data curation C.H., C.Z. and S.C.; Formal Analysis C.H.; Funding acquisition D.S.; Project administration C.H.; Supervision D.S.; Visualization C.H, C.Z.; Writing – original draft C.H., S.C., C.Z., S.R., E.T. and D.S.

## Acknowledgments

We warmly thank all the children and their families as well as Anna Montagnini for her contribution to dataacquisition. Research supported by grants ANR-21-CE28-0010 (DS), ANR-16-CONV-0002 (ILCB) and the Excellence Initiative of Aix-Marseille University (A^*^MIDEX).

## Author Note

In accordance with the Peer Reviewers’ Openness Initiative (https://opennessinitiative.org, Morey, Chambers, Etchells, Harris, Hoekstra, Lakens, et al., 2016), all materials and scripts associated with this manuscript will be available on https://osf.io following publication

